# Structural basis of self-loading by replicative helicases

**DOI:** 10.64898/2026.10.17.759575

**Authors:** Cuncun Qiao, Ignacio Mir-Sanchis

## Abstract

DNA replication is conserved in all forms of life on earth, from subcellular structures such as viruses to human cells. Here we study a group of virus parasites whose replication is initiated by helicases like those found in simian viruses and eukaryotic cells. The process by which these viral helicases initiate the replication of the DNA remains poorly understood. Here, we show how the helicase binds the parasite’s origin of replication with its carboxy terminal domains (CTDs) adopting flexible and different conformations. The CTD’s flexibility moves the DNA towards the inside of the helicase’s ring and bends the DNA at three discrete positions curving it. Our cryo-EM snapshots show that the helicase deforms the DNA while self-loading onto it as the first stage of the initiation of DNA replication, suggesting that initial melting and self-loading are coupled processes in this type of virus parasites.

## Introduction

Initiators of replication are conserved proteins in all domains of life. The main function of initiators is to mark the precise loci where the replication will start by remodeling the duplex DNA so that is unwound generating a small segment of single stranded DNA called a replication bubble ^1^. The replication bubble is the three-dimensional space where several replicative proteins will gather and associate together to form a pre-priming or pre-replication complex ^1–3^. One of those proteins, the replicative helicases, are also conserved in all three domains and always dispose a characteristic architecture as hexameric rings ^4^. To perform their helicase function during replication, replicative helicases need to be loaded onto the DNA so that in their final functionally active state, one DNA strand threads the hexameric ring while the other strand gets excluded: referred to as the ‘replicative mode’. The loading process is usually aided by helicase loader proteins ^5^. There are examples where replicative helicases carry a self-DNA binding domain so that helicase loaders are not needed, thus self-loading onto the DNA. Self-loading helicases are common in viral genomes and associated parasites ^6–9^.

Both initiators and replicative helicases bind adenosine triphosphate (ATP) with a dedicated adenosine triphosphatase domain (ATPase) which classifies them within the large family of proteins with diverse cellular activities AAA+ ^4^. As other AAA+ proteins, initiator proteins use their ATPase domain to form large multi-subunits complexes. The ATP is bound and hydrolyzed and the boundaries between subunits. Such macromolecular architecture allows initiators and helicases to remodel their target DNA molecule. In addition to the ATPase domain, initiators recognize and bind the target DNA by using a DNA binding domain composed of a winged helix motif (WH), which is a common motif in DNA binding proteins. Two elements form the winged helix motif; a helix turn helix and a beta hairpin. Among the three domains of life, DnaA is the initiator in E. coli; Orc in archaeal and ORC in eukaryotes. Despite divergent sequences, they employ similar spatial disposition of their domains to achieve their ultimate goal of replication initiation ^10–12^. Several structures of initiators and replicative helicases from cellular organisms and from viruses have been solved and provide a robust body of knowledge about their functions ^7,13–19^. However, the mechanism by which self-loading replicative helicases end up loaded onto the DNA remains poorly understood.

Eukaryotic viruses, bacteriophages and their parasites represent the major reservoir of genetic content in the planet ^20,21^. To maintain such immense amount of genetic material, they have evolved a high variety of mechanisms to replicate their DNA ^22^. The prototypical family of bacteriophage parasites, the Staphylococcus aureus Pathogenicity Islands (SaPIs), are the focus of this work ^6,23^. Throughout the manuscript we refer to them as SaPIs or islands interchangeably. Different SaPIs carry different accessory genes, generally linked to pathogenesis. This way, SaPI1 and SaPI2, the first islands whose mobility was experimentally demonstrated, carry the *tst*, the gene for toxic shock syndrome toxin-1 ^24^. The SaPIs mimic the genetic organization seen in bacteriophages, where genes with related functions are clustered together in modules ^6,25,26^. Replication wise, these islands encode three elements that allow them autonomous replication: a primase, an initiator of replication, and the origin of replication (*ori* DNA) always located downstream to the primase-initiator genes ^27,28^. The initiators of replication are called Rep. Sometimes the *pri* and *rep* genes are fused, thus generating a PriRep protein. Reps also provide the helicase function during the replication of the islands. For simplicity we refer to Reps as helicases and not as initiators, since they fold as the canonical overall architecture of prokaryotic replicative helicases as homo hexameric rings ^29^. In its replicative mode, translocation along the DNA separates both strands which is a necessary condition to provide the DNA template to the DNA polymerase ^30^. Here Reps fold as a three-tiers rings. Their amino terminal domain has undefined functions. The central domain is the ATPase, upon which we use to classify Reps in three prototypes based on structural homology. Rep 1, Rep 2 and PriRep 5 encoded by SaPI1, SaPI2 and SaPI5 respectively (Supplementary Figure 1). Rep 1 and PriRep 5 fall into the Super Family 3 (SF3) of helicases and Rep 2 as an archaeal and eukaryotic MCM homolog in the SF6 of helicases. The third tier is the CTD whose winged helix motif is used to bind to their cognate *ori* DNA while acting as initiators of replication ^27^. Since there are not accessory factors loading Reps onto the *ori* DNA, they are considered self-loading helicases. The 300 bp-spanning *ori* DNA shows a conserved structure of discrete penta- or hexa-nucleotides called iterons which are oppositely disposed surrounding an AT-rich region of DNA (Supplementary Figure 1). The AT-rich region is assumed to be melted upon initiation. The Rep’s CTD binds specifically to the iterons, presumably one hexamer on each side of the AT-rich region. Structures of Rep hexamers and their homologs have been solved in apo state and bound to single stranded DNA (ssDNA), the aforementioned replicative mode ^7,8^. However, structures of these machines in complex with *ori* DNA acting as initiators are still missing, limiting our current understanding on how these ringed helicases self-load onto and melt the DNA. Here we present two different snapshots of PriRep5 in complex with *ori* DNA to reveal an unseen capacity to bind and bend the DNA during the self-loading process. PriRep5 maintains its hexameric quaternary disposition as a barrel, although the hexamer is expanded compared to the replicative form. The first structure captures four CTDs contacting the *ori* DNA, one of which swings out to bind the iteron that happens to be in an inverted orientation. The second snapshot shows three ordered CTDs and a robust distortion of a segment of DNA that is not the canonical AT-rich region but that is compatible with an initial melting of the segment that is unexpectedly AT-rich. Our data suggest that initial melting of DNA occurs concomitantly to the self-loading process. This work widens the current knowledge on *ori* recognition and helicase self-loading mechanism onto DNA during the initiation of replication of these islands.

## Results

### Cryo-EM structure of PriRep5 bound to its ori DNA

PriRep5 was incubated in complex with partial origin DNA comprising 105 base pairs and MgCl_2_ (see methods, supplementary figure 1). The 105 bp oligonucleotide corresponds to the upstream region of the ori, excluding the canonical AT-rich region (green) and the set of iterons in the downstream region. Within the iterons included in the oligonucleotide, iterons L1 and L5 show an inverted disposition respect iterons L2,L3,L4 and L6 (Supplementary Figure 1C). We selected this upstream region because here the iterons seem to have one more conserved nucleotide in their sequence, TGCAACC, as opposed to TGCAAC on the other downstream side of the *ori*. We collected 3412 movies and generated a consensus cryo-EM map at an overall resolution of 4.2 Å (methods, Supplementary figure 2). Although in PriRep5 the primase is fused to the helicase, the primase domain remains invisible in the map, which is a common feature in these types of helicases ^8^. What is evidenced in the structure is PriRep5’s characteristic hexameric barrel architecture with three CTDs ordered, which are contacting a long density corresponding to a double stranded DNA (Figure 1). Focused refinement of the CTDs-DNA region generated a better map where four CTDs and a longer segment of DNA were visible (Figure 1). The local resolution of this focused map ranges from 3.2 to 5 Å in most part of the DNA (Supplementary figure 2). Notably, the major and minor groves of the DNA are clearly identifiable along the whole density. We call this structure PriRep5-4CTDs. The overall shape of PriRep5-4CTDs is equivalent to that seen in the replicative mode with the particularity that here the hexamer seems to be breathing and consequently it is expanded, slightly moving apart adjacent ATPase subunits (Supplementary figure 3A). The distance between the outer amino acid residues G471 of opposite subunits in the replicative mode was 111.174 Å, 111.349 Å, and 111.991 Å for subunits A-D, B-E and C-F respectively, compared to 125.585 Å, 115.839 Å and 117.608 Å in PriRep5-4CTDs. This might be the reason that explains why PriRep5 does not need ATP to bind and melt its *ori* DNA ^27,28^. The conformational disposition of the barrel to bind the ori DNA seems incompatible with nucleotide accommodation between subunits. The four CTDs make clear contact with the long density corresponding to 62 base pairs of the origin DNA with one extra nucleotide in strand assigned as chain G. The main component of PriRep5’s CTDs is the WH commonly seen in DNA binding proteins. Here, the winged element is the antiparallel beta sheet composed by beta strands #3 and #4 (Figure 2 and Supplementary Figure 4). Contrary to other DNA binding proteins using the ‘wings’ of the winged helix turn helix motif to bind the minor grove of the DNA^11,31^, PriRep5 uses its ‘wing’ to land on the major grove and interact with the bases, suggesting that the antiparallel beta sheet is the DNA binding element responsible for iteron sequence recognition. Additionally, the helix #4 of the WH binds the minor grove’s backbone. Interestingly, the CTD of the subunit we named arbitrarily as A has swinged out extensively and binds the DNA from the side instead of binding it from the top (Figures 1 and 2 and Supplementary figure 3B). Still, despite the rotation of subunit A’s CTD, it binds the DNA in a similar fashion as the other CTDs: the helix turn helix binds the minor grove’s phosphate backbone and the winged beta sheet binds the bases in the major grove, but this time of the complementary strand (Figure 2). We hypothesized that this was the consequence of the iterons’ disposition and built the sequence of the DNA model trying to align the sequences of the iterons with the CTDs. Indeed, they nicely align and subunit A’s CTD (CTD-A) binds iteron L5 while CTD-B, CTD-C and CTD-D bind to iterons L4,L3 and L2 respectively. (Figure 2 and Supplementary Figure 1).

**Figure 1.**
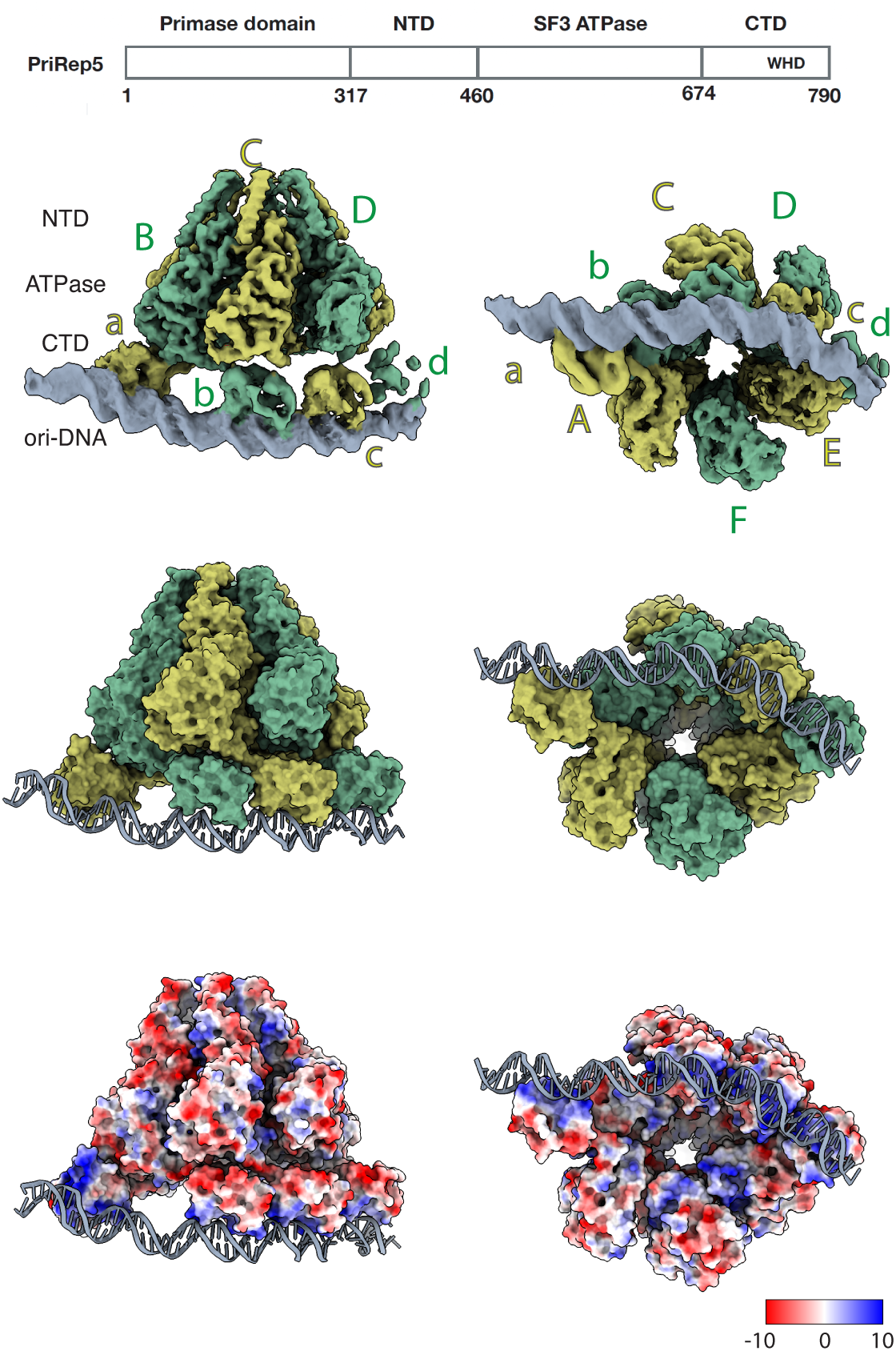
Structure of PriRep5-4CTDs. Top, scheme of PriRep5 polypeptide chain. The numbers represent amino acid residues. NTD, amino terminal domain, SF3 ATPase, superfamily 3 ATPase domain, CTD, carboxy terminal domain, WHD, winged helix domain. Middle top, orthogonal views (left: lateral view; right: bottom view) of the cryoEM map of PriRep5-4CTDs. Subunits are colored gold and green, while ori DNA is colored grey. NTD and ATPase domains are labelled with capital letters and CTDs are labelled with lower case to evidence that CTD of a subunit is positioned underneath the adjacent subunit. Middle Bottom, two orthogonal views of PriRep5-4CTDs model shown as surface representation. Same colors as in the middle panel. Bottom, PriRep5 surface representation as electrostatics and the *ori* DNA as grey cartoon.

**Figure 2.**
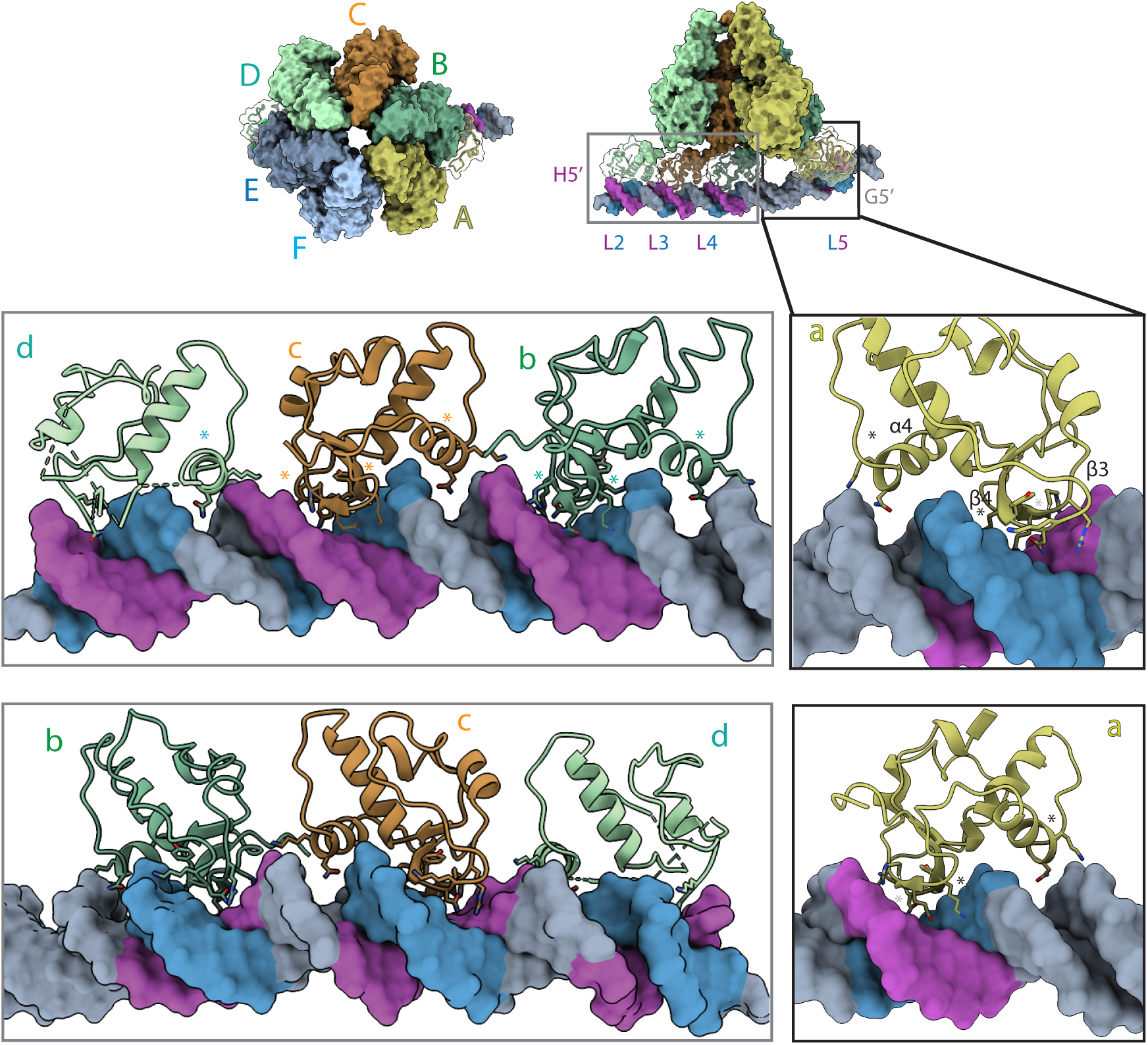
Ori recognition by PriRep5. Top, ring top view of PriRep5-4CTDs where the protein occludes the DNA almost entirely. Chains A to F are colored yellow, dark green, orange, light green, dark blue and light grey respectively and shown as surface. DNA is shown as surface and colored both strands dark grey, with iteron sequences 5’-TGCAACC-3’ as purple. In the orthogonal view shown on the right (lateral view), subunits E and F are removed for clarity. The CTDs are shown as ribbon, and the DNA becomes more visible where iterons with sequence 5’-GGTTGCA-3’ are now evidenced as blue. Chains G and H correspond to the DNA strands. Labels G5’ and H5’ are positioned on the corresponding 5’ end. The label H5’ is colored purple to help identify the strand, which is the one on the top, starting with the colored purple sequence 5’-TGCAACC. L2, L3,L4 and L5 represent the iterons as labeled in supplementary figure 1. Boxed regions have been zoomed in to see the CTDs in contact with the DNA. The boxes on the bottom have been flipped 180 degrees respect the vertical axis to show the contacts from the other side. Side chains from residues N729 and K730G (α4); Q748, R749 and R750 (β3); and K777, N778, Y779 and K780 (β4) are shown as sticks. Stars mark α4, β3 and β4. See methods and main text.

### PriRep5 bends and moves the DNA

The structure of PriRep5-4CTDs shows how the helicase bends and distorts the DNA in several manners. When looking at the structure from the bottom, the DNA is bended at three different positions, generating four segments of DNA with different long axis directionality (Figure 3). Because of such bending the DNA appears with a certain curvature. When looking at the structure from the side, the DNA is bended in a different direction and at two different points. Notably, the region of the DNA between CTD-A and CTD-B is the one that suffers those bending events in different directions. We analyzed the DNA deformation by performing three types of measurements compared to the canonical B-form of DNA: backbone deformation as the phosphate-phosphate distance between complementary nucleotides; DNA grove deformation as measured by phosphate distances; distance between donor-acceptor atoms in Watson-Crick base pairs hydrogen bonding as an indication of DNA melting. The measure of backbone deformation between phosphates of complementary pairs (Supplementary Figure 5) indicated that out of 27 nucleotides analyzed in iteron sequences, 14 showed increase distance while 9 showed decreased distance with the remaining 4 nucleotides exhibiting the same distance as the B-form. The DNA grove measurement did not evidence any identifiable deformation pattern in the iteron sequence. It should be noted that the script used in the measurements would detect winding, unwinding and bending deformation, thus all deformations contributing to a different width measurement in DNA grove. The Watson-Crick base pair distances measurement showed that the DNA captured in the complex is not being melted, as all distances were like those present in the canonical B-form of double stranded DNA (Figure 3 C).

**Figure 3.**
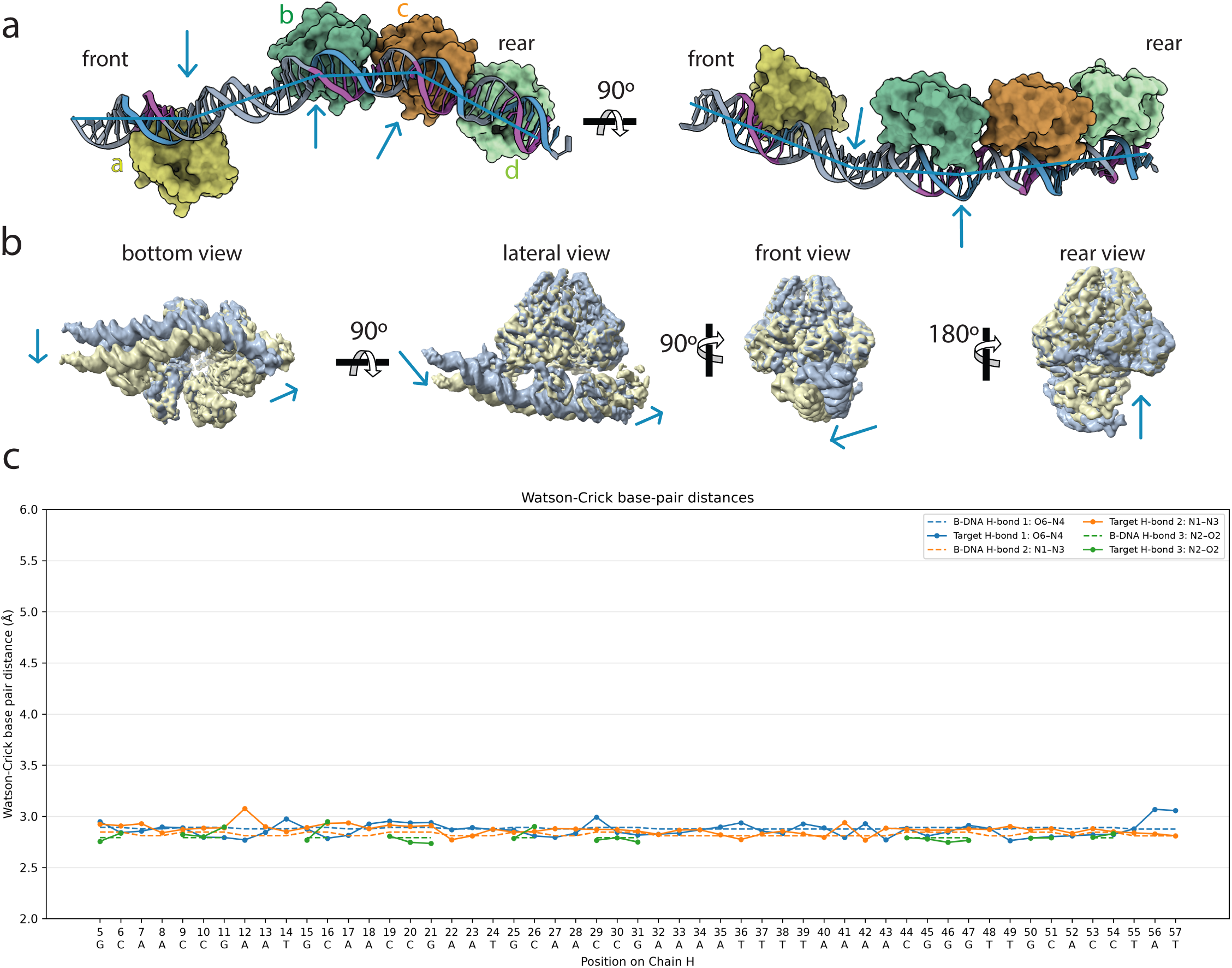
DNA bending and movement. A. Orthogonal views (left bottom view, right lateral view) of PriRep5-4CTDs where the NTD and ATPase domains of all subunits have been removed for clarity. The CTD-A, CTD-B,CTD-C and CTD-D are shown as surface and colored yellow, green, orange and light green respectively. The DNA is shown as ribbon where the helix axis is highlighted with a blue line to evidence the DNA bending. The iteron sequences 5’-TGCAACC-3’ and 5’-GGTTGCA-3’ are colored as in Figure 2 purple and blue respectively. The blue arrows indicate the points at which the DNA is bent. B. DNA movement analysis. The two more distant intermediate maps generated during multibody refinement are shown as blue and pale-yellow surfaces. The blue arrows indicate the movement from the blue map to the pale-yellow map. See supplementary figure 5 and the supplementary videos for a complete movement by the ten intermediates. C, Bottom panel, plot of atomic distances between Watson-Crick base pairs compared to canonical DNA in its B-form. The atomic distances are measured between atoms O6-N4 (or N6-O4), N1-N3, O2-N2 and plotted as H-Bond 1 , 2 and 3 respectively. The distances in the canonical B-DNA are shown as discontinued lines, while the distances measured in PriRep5-4CTDs are shown as connected dots. The horizontal axis shows the identity and number position of chain H nucleotides.

We hypothesized that the complex captured in our dataset is not only binding the origin of replication but is also initiating helicase loading onto the DNA. To test this hypothesis, we analyzed the flexibility of the DNA-bound helicase’s CTDs by performing variability component analysis by running multibody refinement in RELION ^32,33^. As shown in Figure 3B, supplementary figure 6 and supplementary videos, the DNA-bound helicase showed a set of different intermediate states performing the movement of the DNA towards the center of the ring, indicating that the loading mechanism was initiated and captured in our dataset.

Further processing of our dataset allowed us to generate another map that we called PriRep5-DNA-distorted (Figure 4, Supplementary Figure 2). In this map, three CTDs were ordered. Interestingly, the DNA visible in this intermediate was shorter and it showed a stronger degree of deformation. Not only in the regions where the wings interact with the DNA but also in the region between CTD-A and CTD-B. At a contour level of 0.00913, where the map shows continuous density for the totality of the complex, this region of the DNA shows how both strands of DNA are being separated (Figure 4A). The local resolution for this map in this region reaches 3.5 Å resolution (Supplementary figure 2). As with the previous map, the WH motif employs the helix component to interact with the minor grove’s backbone and the wing component for major grove interaction and iteron recognition (Figure 4A). We analyzed the deformation of this model’s DNA (Figure 4 B and Supplementary Figure 5). In this case, the Watson-Crick base pair distances showed a considerable separation in nucleotides C16, C20, A23, C26, T38 and G50. Notably, the affected cytosines fall in L3, L4 and L5 iterons, while the adenine and the thymine fall in the flanking regions between iterons. We hypothesized that if the region between iterons was indeed a melting point, it should be AT-rich. We then analyzed the sequence between iterons and discovered that iterons L2, L3 and L4 were separated by the dinucleotide AA (chain H, Supplementary Figure 1), while the sequence between L4 and L5 was indeed AT-rich: out of 15 base pairs, 12 were formed by A-T tracks of four bases alternating A and T in the form of AAAATTTTAAAA. We then searched for a similar structure on the other side of the ori: inverted iterons separated by 17 AT rich base pairs. We searched for 17 nucleotides instead of 15 because the iterons in the downstream region have one less nucleotide conserved. We found that iterons R2 and R3 in the downstream region which are 17 base pairs apart, are separated by a sequence containing the AT rich bit AAAATTTAAAA (Supplementary Figures 1). We then searched for such structure in the other two prototypical islands SaPI1 (SF3 helicase Rep1) and SaPI2 (SF6 MCM-like helicase Rep2). In both cases we could identify iterons and sequences compatible with the hypothesis that such locus might represent initial melting points (Supplementary Figure 4). Moreover, our data agrees with previous experimental data where deletion of this set of iterons and the segment between inverted iterons prevents *ori* melting and DNA replication of SaPIbov1 island *in vivo* ^28^. To confirm our structural data and demonstrate that the WH is responsible for the DNA interaction, we performed structure-based mutagenesis of the amino acid residues present in α-helix 4, β-strand 3 and β-strand 4 (Supplementary Figure 4). We generated three sets of mutations that we named α4, β3 and β4. Mutation α4 bears N729A and K730G; mutation β3 carries Q748A, R749A and R750A, while β4 does so with K777A, N778A, Y779A and K780A. Electromobility shift assays (EMSAs) demonstrated that β3 and β4 mutants completely abolished their DNA binding activity, while mutant α4 was compromised as compared with the wild type but it did not abolish its function as it was the case for the β mutants.

**Figure 4.**
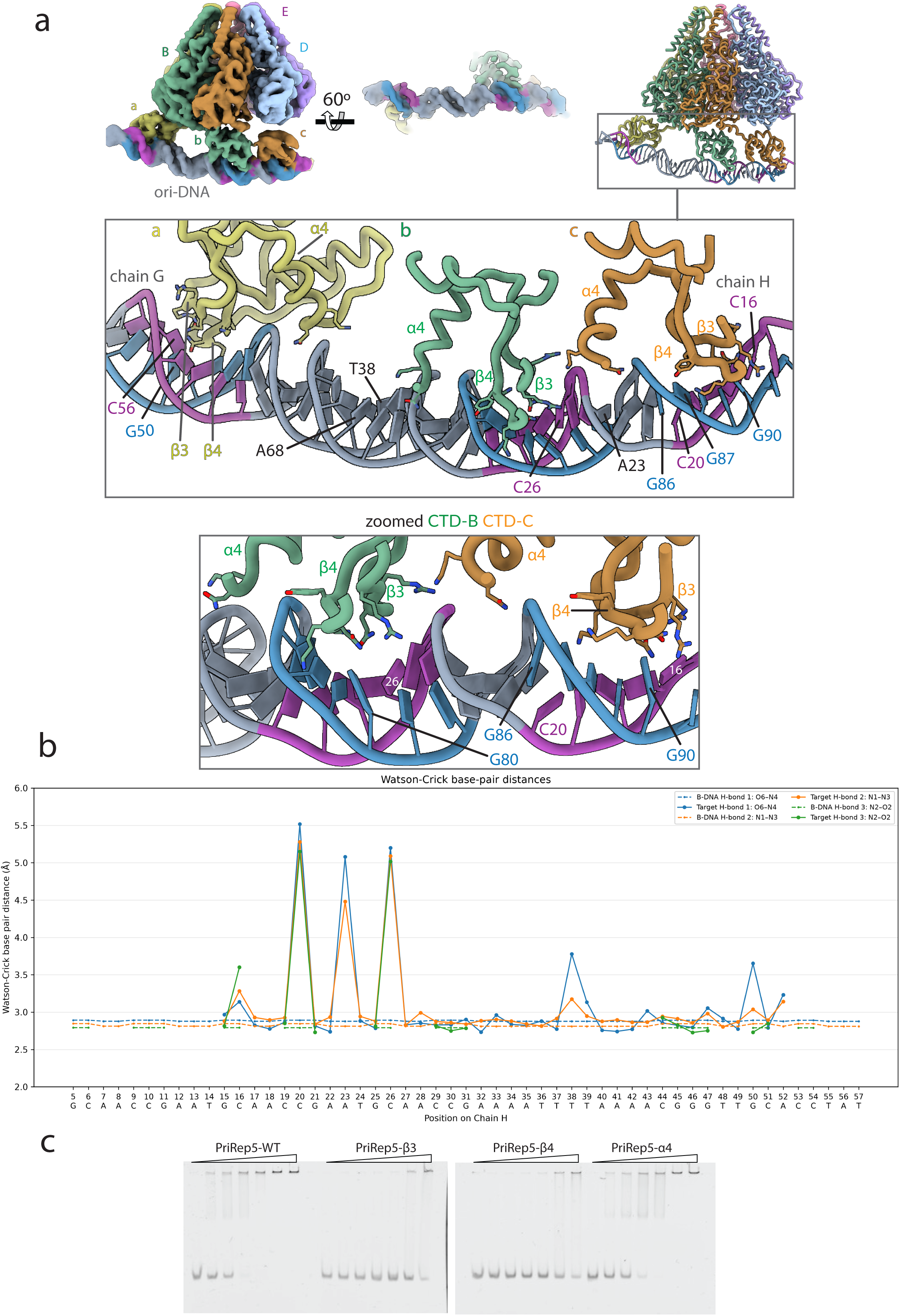
ori DNA deformation by PriRep5. A. Cryo-EM map of PriRep5-DNA-distorted colored as chain colors in the cartoon shown on the right as ribbon representation. Inlet of a zoomed in vision of CTD-A and CTD-B contacting the *ori* DNA. Mutated residues N729 and K730G (α4); Q748, R749 and R750 (β3); and K777, N778, Y779 and K780 (β4) are shown as sticks and highlighted in supplementary figure 4. Within CTD-B (green), the residues 704-728, 752-775 are removed for clarity. B, Plot of atomic distances between Watson-Crick base pairs compared to canonical DNA in its B-form. The atomic distances are measured between atoms O6-N4 (or N6-O4), N1-N3, O2-N2 and plotted as H-Bond 1 , 2 and 3 respectively. The distances in the canonical B-DNA are shown as discontinued lines, while the distances measured in PriRep5-ori-distorted are shown as connected dots. The horizontal axis shows the identity and number position of chain H nucleotides. C, EMSAs of PriRep5 wild type and the mutants α4, β3 and β4 using *ori* DNA PCR amplified as substrate. Protein concentrations are 0,2,5,10,20,50 and 100 nM. See methods.

## Discussion

The current structural information about SaPI’s and other staphylococcal genomic islands’s initiator/helicases mainly focuses on their replicative mode ^7,8,34^. We provide here structural understanding of how the self-loading replicative helicase PriRep5 from SaPI5 recognizes its origin and starts the initiation step during SaPI replication. PriRep5 performs quite considerable conformational movements to achieve origin recognition as compared to the replicative mode. First, the hexameric ring is expanded as evidenced by the ATPase tier, which shows displacement between subunits to accommodate the extensive opening of the CTD tier. Second, two types of CTD displacements are evident: those performed by CTD-B,CTD-C and CTD-D; and that seen in CTD-A which represent the most drastic conformational change to accommodate binding to inverted iteron #5. As opposed to archaeal Orc initiators, where the winged element inserts into the minor grove of the origin DNA ^11,12^, PriRep5 positions the winged element of its WH in the major grove, employing Q748, R749, R750, K777, N778, Y779 and K780 as critical residues in *ori* DNA recognition. Another difference with Orc initiators is that here, the ATPase domain does not interact with the DNA whatsoever. We should not rule out the possibility that the ATPase-DNA interaction occurs in a subsequent intermediate state not captured in our data set. One feature we could identify in our data set was the ample movements that DNA-bound CTDs exercise. Although CTD flexibility was identified earlier for the apo state of Rep1 and PriRep5 ^8^, it was uncertain if such flexibility was maintained in a DNA-bound intermediate. The variability analysis we performed here confirms that DNA-bound CTDs are highly flexible and that the CTDs can approximate the DNA towards the ring. Moreover, our structural analysis showed that within our data set we could identify the movement of the DNA towards the ring and ori DNA distortion which is compatible with *ori*-DNA melting. Our in-silico analysis discovered that the sequence between iterons is as AT-rich as the canonical AT-rich region (Supplementary Figure 4). We do not claim that the canonical 102 bp long AT-rich region in the center of the SaPI5 origin is not used for melting, since deletion of this region in SaPIbov1 prevented origin melting and island’s replication *in vivo* ^28^. Nonetheless, our data suggest that there are several melting points in addition to the large central AT-rich region, which agrees with experimental evidence that showed that flanking iterons plus the sequence between inverted iterons are a necessary condition for a fully active origin of replication *in vivo* ^28^.

Our study’s limitations prevent to clarify how two hexamers fully associate with the island’s origin DNA. Similarly, important intermediates are still a mystery. The precise motions and conformational changes that allow the hexamers to end up fully loaded onto the DNA with a fully open replication bubble are still unknown. Such information will be fundamental to continue analyzing conserved, converged and diverged strategies among replicative systems.

## Methods

### Plasmid cloning and PriRep5 protein expression and purification

The lists of oligonucleotides and plasmids are found in supplementary tables 2 and 3 respectively. DNA encoding the wild-type SaPI5-encoded prirep5 gene was amplified by PCR from the USA300 genome. The PCR product was cloned into plasmid pET28a using the restriction enzymes NcoI/XhoI, respectively generating the plasmid pIMS520. For site-directed mutagenesis, the plasmid pIMS520 was used as template to amplify the desired mutations alpha4, beta3 and beta4, with oligonucleotide pairs listed in supplementary table 2. The resulting PCR products were digested with 1 U DpnI at 37 ℃ for 1 h and then transformed into E. coli DH5α. The sequence confirmed clones were transformed into BL21 (DE3) Rosetta cells which were grown in LB medium supplemented with 50 µg/ml kanamycin and 25 µg/ml chloramphenicol at 37 ℃ until they reached an OD600 of 0.6–0.8. Protein expression was induced by addition of 0.5 mM IPTG at 18℃ for 16 h. Cell pellets were harvested and re-suspended in Buffer A1 containing 50 mM sodium phosphate (pH 7.0), 5% glycerol, 1 mM DTT, and 1 M NaCl in the presence of a protease inhibitor cocktail (Roche) and then lysed with 0.5 mg/mL of lysozyme for 30 min at 37℃ followed by sonication. After centrifugation, the supernatant was loaded onto a HisTrap^TM^ HP column equilibrated with Buffer A1, and the targeted protein was eluted with a gradient from 0 mM to 500 mM imidazole using an ÄKTA pure system. Elutions were concentrated and injected into HiTrap Heparin HP column (Cytiva) equilibrated with the appropriate A2 buffer and eluted with a NaCl gradient up to 2 M. Peak fractions were pooled, concentrated and injected into HiLoad® 26/600 Superdex® 200 pg columns (GE Healthscience) equilibrated with Buffer B (20 mM Tris pH 8, 200 mM NaCl, 0.5 mM EDTA, 2 mM DTT). Fractions were analyzed by SDS-PAGE, and those fractions containing the target protein were pooled, concentrated and dialyzed with stock buffer (20 mM Tris pH 8, 200 mM NaCl, 0.5 mM EDTA, 20% glycerol, 2 mM DTT). After dialysis at 4°C, the preparations were flash-frozen in liquid nitrogen and stored at –80 °C.

### Electrophoretic mobility shift assay (EMSA)

FAM-labeled oligonucleotide ori5-TET-F and Ori_SaPI5_HindIII_R were used to amplify SaPI5 origin of replication from strain USA300. 2 nM of substrate dsDNA was mixed with PriRep5 protein in binding buffer (20 mM Tris, pH 8.0, 100 mM NaCl, 5% glycerol, 50 ng/µl BSA, 5 mM DTT, 10 mM MgSO_4_, and 1 ng/µl salmon sperm DNA). The total volume of the reaction was 20 ml. After incubation at 37°C for 30 min, 4 µl of loading buffer (20 mM Tris pH 8.0, 60% glycerol, 30mM EDTA and 0.03% xylene cyanol/bromophenol blue) was added to the reaction and the entire volume was then loaded on native 4% polyacrylamide gels. Gels were run in 0.5× TBE at 4°C for 2 hours. The gels were imaged using a Typhoon 9400 (FLE).

### Specimen preparation for Cryo-EM and data collection

Oligonucleotides ori_105nt_SaPI5 and ori_105nt_SaPI5-R were annealed as previously described in Qiao et al, 2022). 1 mg/ml of PriRep5 was incubated with approximately 2-fold excess of duplex oligonucleotide and 5 mM MgCl_2_ for 30 min at 4°C in binding buffer. 4ul of sample was applied to freshly glow discharged Quantifoil 2/1 carbon grids, blotted for 4 s, and flash frozen in liquid ethane using a Vitrobot (Thermo Fisher Scientific) with a controlled humidity of 100% and temperature of 4°C. The dataset was collected using a Glacios Cryo Transmission Electron Microscope operated at 200 kV with a 100 µm objective aperture and a nominal magnification and pixel size of 150,000x and 0.953 Å per pixel respectively. Complete data collection parameters are summarized in supplementary table 1.

### Image processing and 3D reconstruction

The workflow for cryo-EM data processing is illustrated in Supplementary Figure 5. Initial data processing was carried out in the CryoSPARC 4.2.1 Live environment ^35^, where 3412 raw images were motion-corrected and CTF-estimated on-the-fly during data collection and then filtered based on CTF fit estimation and relative ice thickness. Blob picker was first used to pick the particles in 200 micrographs. The particles were then used for generating several 2D averages which were subsequently used as a template for template picking. Template picking was then performed for the whole dataset, and 3 586 153 particles were picked. Two template pick jobs were maintained in parallel to be subjected to five and four rounds of 2D classification, after which a total of 416,918 particles were picked (186,876 and 230,042 particles respectively). After verification that those sets of particles generated good maps in CryoSPARC ^35^, the coordinates of those particles were used in RELION 5.0.1 to extract 2xbinned particles from micrographs which had been imported, motion corrected ^36^ and CTF calculated ^37^ previously in RELION. For motion correction we used RELION’s own built-in CPU implementation. Subsequent initial models, 2D and 3D classifications, 3D and focused refinement jobs were conducted in RELION. Local resolution maps were generated by running ResMap ^38^ as implemented within the RELION wrapper interface.

For PriRep5-4CTDs, the initial model was generated by imposing C1 symmetry to the set of 186,876 particles, then 3D refined followed by 2 rounds of 3D classification, where a total of 112,448 particles were then joined to the other set of particles to run 3 rounds of 3D classification where one class yielded 49167 particles that were re-extracted unbinned. The consensus map at 4.2 Å was generated after one round of 3D, polish and CTF refinements. Such map showed density for 3 CTDs and for *ori* DNA. A particle subtraction job encompassing the CTD regions of PriRep5 plus the *ori* DNA preceded a focused refinement that generated the Focused map at 4.9 Å overall resolution where 4 CTDs were visible and substantial larger density corresponded to the DNA. In such density, as in the consensus map, the DNA major and minor groves where perfectly identifiable.

For PriRep5-DNA-distorted the 49167 particles were used to generate an initial model imposing C6 symmetry followed by two 3D refine jobs with and without symmetry that yielded a consensus map at 4.6 Å overall resolution. Here the particle subtraction and focused refinements involved both the hexamer domain (helicase) and the CTD-DNA region, that generated a focused map of 3.5 and 4.4 Å overall resolution respectively. The mask for the CTD-DNA region included a large region of the ATPase domain of subunits A and B that is highlighted in supplementary figure 2 data processing workflow.

For modeling, AlphaFold3’s ^39^ prediction of the hexamer was used to fit in the map followed by several rounds of automatic flexible fitting with Namdinator ^40^, manually corrected in Coot ^41^ and real space refinement in Phenix ^42^. As a final step, the model was refined with Isolde ^43^ using distances and torsions restrains from the Alphafold3 model as template.

### Atomic distances measurement in the DNA

To analyze the atomic distances in the DNA atoms in both PriRep5-4CTDs and PriRep5-DNA-distorted, first a template double stranded DNA with the same sequence as the duplex oligonucleotide used in the specimen preparation was created in Coot as an ideal B-form of double stranded DNA. Then three different python scripts were generated to measure atomic distance between phosphate atoms of complementary base pairs [analyze_dna_pp.py], between phosphate atoms of nucleotide residues separated a fixed distance before and after the target nucleotide [analyze_dna_grooves.py], and between donor-acceptor atoms of the Watson-Crick pairs [analyze_dna_hbonds.py] in both the template B-form DNA and the target, either PriRep5-4CTDs or PriRep5-ori-DNA. Given the numbering assigned to the nucleotides in chains H and G, for the grove measurement of a particular n residue in chain H (H:n), the major groove is measured between H:n and G:(101-n). For the minor groove, the distance is measured between H:n and G:(111-n). This way, for H:45, the major groove is measured with G:56, while the minor groove is measured with G:66. The scripts log the values in a CSV file that is included as a supplementary data file called measurements. The plots represent the values from both the template B-form and the target. Scripts are available upon request. The analysis of atomic distances in PriRep5-4CTDs extended from chain H nucleotide 5 to nucleotide 57, and from 17 to 52 in PriRep5-ori-DNA-distorted due to the different length of DNA density present in each structure.

Software used in this work was curated by SBGrid ^44^.

## Data availability

Coordinates and volumes are deposited in PDB and EMDB under the following accession codes: PriRep5-4CTDs coordinates PDB ID 32XM; PriRep5-4CTDs Composite map EMD-59245; PriRep5-4CTDs Consensus map EMD-59247; PriRep5-4CTDs Focused map EMD-59248; PriRep5-DNA-distorted coordinates PDBID 32DE; PriRep5-DNA-distorted Composite map EMD-59749; PriRep5-DNA-distorted Consensus map EMD-58826; PriRep5-DNA-distorted Focused Map helicase domain EMD-58827; PriRep5-DNA-distorted Focused Map CTD domain EMD-58829.

## Supporting information

Supplementary Figure 2 -FSC

## Acknowledgments

We thank the Knut and Alice Wallenberg Foundation, members of the Wallenberg Centre for Molecular Medicine Umeå, Umeå University and Region Västerbotten. We thank Michael Hall and Camilla Holmlund for their help in specimen preparation, screening, and data collection at the Umeå Core Facility for Electron Microscopy, a node of the Cryo-EM Swedish National Facility, funded by the Knut and Alice Wallenberg, Family Erling Persson and Kempe Foundations, SciLifeLab, Stockholm University and Umeå University. Initial computations were enabled by resources provided by the Swedish National Infrastructure for Computing (SNIC) at HPC2N partially funded by the Swedish Research Council through grant agreement no. 2018-05973. This work was co-funded by the Swedish Research Council under grant number 2023-03117 to I.M-S, by the Spanish Ministerio de Ciencia, Innovación y Universidades (MICIU) and the Agencia Estatal de Investigación (AEI) under Grants PID2024-158622OB-100 and RYC2022-035465-I to I.M-S.

## Authors Contributions

C.Q., methodology, data curation, investigation, formal analysis, resources, validation, writing original draft. I.M-S., conceptualization, validation, resources, supervision, writing.

---

**Supplementary Figure 1. Genomic context of SaPI’s origin of replication**. A, annotations for Rep1, Rep2/Repbov1 and PriRep5. B, arrows indicated primase and replicase genes with the origin of replication downstream the rep gene. The inlets show the iteron’s sequence and the number of repeats. US, upstream, DS, downstream respect the central AT-rich region. C, sequence of SaPI5 origin of replication. The last 12 residues of the *prirep* gene are highlighted along with the iterons, which are labeled as L (left)1 to 6 and R (right) 5 to 1. The inverted iterons are colored orange. The AT-rich region is highlighted in green. D, sequence of the duplex oligonucleotide used in the mix with the nucleotide numbers as they were assigned in our PriRep5-4CTDs and PriRep5-DNA-distorted structures. Underline and pink mark the DNA modeled in PriRep5-4CTDs and PriRep5-DNA-distorted respectively.

**Supplementary Figure 2. Cryo-EM processing workflow**. See methods. When the same map is shown twice, those images are the orthogonal views. The FSC curves are shown for the Consensus [EMD-59247] and Focused [EMD-59248] maps regarding PriRep5-4CTDs, and Focused maps of Helicase [EMD-58827] and CTD [EMD-58829] domains regarding PriRep5-DNA-distorted maps. Below the FSC curves, the masks used for the focused refinements of the CTDs-DNA regions are shown as pink (PriRep5-4CTDs) and yellow (PriRep5-DNA-distorted). The boxed region on the yellow mask highlights the ATPase region included in the mask that is not included in the pink mask. The angular distributions of the Consensus maps are shown on the right-hand side. The two images are rotated 180 degrees in the vertical axis.

**Supplementary Figure 3. PriRep5-4CTDs hexamer is expanded respect its replicative form**. A, Left, a crossed section of PriRep5-4CTDs at its ATPase level is shown as ribbons where subunits A,C,D and B,D,E are colored gold and green respectively. The cartoon is overlapped on its replicative form PDBID 7OLA displayed as pink ribbon. Right, Glycine 471, which is located at the periphery of the hexamer is used to measure the ring diameter as the distance between opposed subunits’ alpha carbons of Gly471. The distances are shown as Å. B. Comparison of CTDs conformations in PriRep5-4CTDs and its replicative form 7OLA. Left is the bottom view where the DNA is seen before as ribbons, and the protein is observed behind as surface representation. Lower case letters represent the CTDs subunits chain identifier. The bottom image is the same cartoon where the NTD and the helicase domains are removed for clarity. The overlapped images show the different conformations, which are labeled as arrows to evidence the opposite movements that subunit a performs compared to subunits b and c.

**Supplementary Figure 4. Secondary structure of CTD.** A, Secondary structure of PriRep5’s CTD. Protein sequence of PriRep5 from residue 610 to residue 790 is shown as one letter code, where underlined residues are those whose numeric position is shown above. Blue helices and pink arrows represent alpha helices and beta strands in the structure and are labeled as A# and B# respectively. Stars mark the residues mutated in alpha helix A4 (α4), and beta strands B3 and B4 (β3 and β4). B, sequence analysis of inverted iterons and the segment between them from SaPI5, SaPIbov1, SaPI2 and SaPI1. The numbers in parenthesis are the AT nucleotides percentage present in the sequence between inverted iterons (boxed) within the upstream, downstream and the canonical AT-rich regions.

**Supplementary Figure 5. DNA deformation measurements.** A, DNA backbone deformation of PriRep5-4CTDS DNA as measured by phosphate-phosphate distance between complementary nucleotides and relative to the distance found in the ideal B-form of DNA. B, DNA grove deformation of PriRep5-4CTDs relative to B-form DNA. The measurements for a particular nucleotide n in chain H are performed against residue in chain G 101 – n. This is, for nucleotide in position 19 in chain H, the major groove is measured from H:19, to G:82. The minor groove is measured with G 111 – n. Therefore, the minor groove is measured between H:19 and G:92. See methods. C and D are the same measurements in PriRep5-DNA-distorted.

**Supplementary Figure 6. DNA movements.** The ten maps generated by multibody refinement are shown in the four same different orientations as in main Figure 3. Front is considered the side where CTD-A is located, while rear is the side where CTD-D is. From the bottom view, where the reader observes the DNA first and the helicase behind, a 90 degrees rotation in the horizontal axis has been applied to get to the lateral view, where CTD-A is seen on the left-hand side and labelled as a. From the lateral view, a 90-degree rotation in the vertical axis is applied so that CTD-A (front side) moves towards the reader. From the front view a 180-degree rotation in the vertical axis gives the final rear view.

**Supplementary Figure 7. Zoomed in region of PriRep5-ori-distorted**. Same image as in main Figure 4A inlet but no residues in subunits C and C are removed for clarity. Mutated residues are show as sticks.

**Supplementary Figure 8. Model to map fit.** PriRep-5-DNA-distorted model fitted into the focused map. Note that the region of the ATPase domain that the map includes corresponding to the A and B subunits has not been fitted with the model in this figure.

