## Supplementary Figure 1 for "Structural basis of self-loading by replicative helicases"

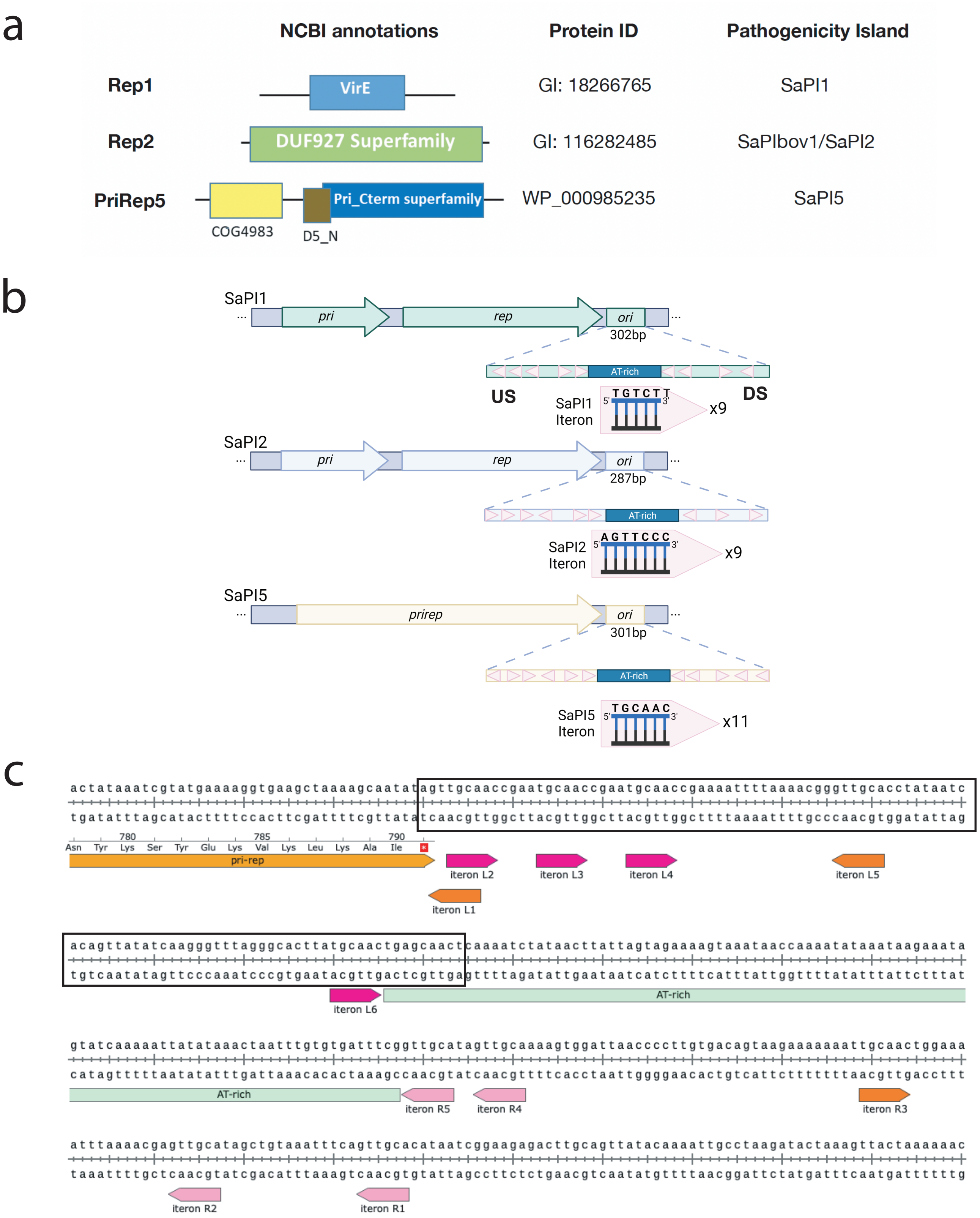


d

```
Chain H
   1  4          14                                          54 57                                              105
5'- agttgcaaccgaatgcaaccgaatgcaaccgaatgcaaccgaaaattttaaaacgggttgcacctataatcacagttatatcaagggtttagggcacttatgcaactgagcaact -3'

3'- tcaacgttggcttacgttggcttacgttggcttttaaaattttgcccaacgtggatattagtgtcaatatagttcccaaatcccgtgaatacgttgactcgttga -5'
      103          92                                          52 49                                               1
                                                                                                                     Chain G
```
