## Supplementary figures and images for "Structural basis of self-loading by replicative helicases"

### Supplementary Figure 2

Supp. Figure: Cryo-EM Processing Workflow

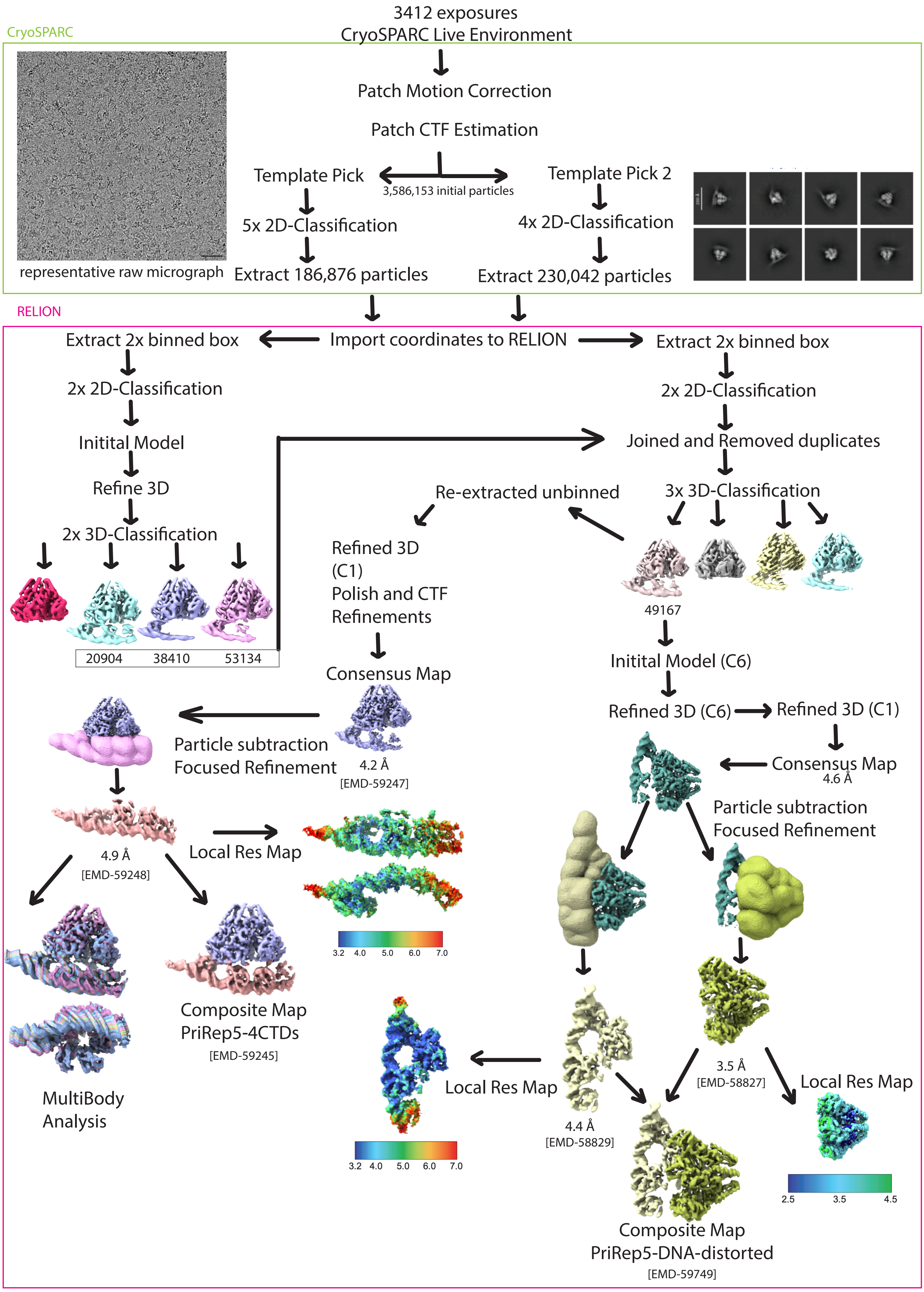

### Supplementary Figure 3

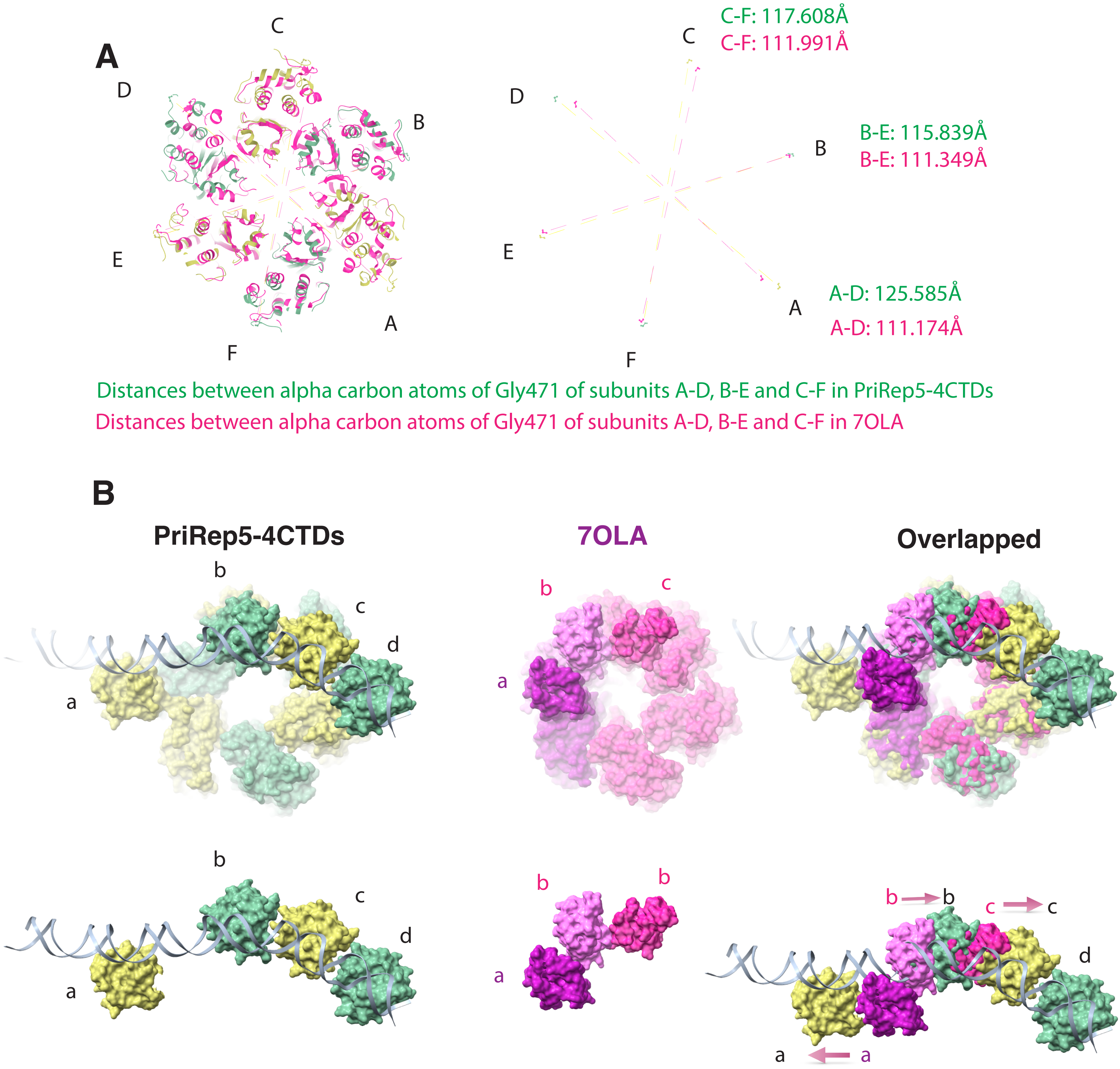

### Supplementary Figure 5

#### PriRep5-4CTDs

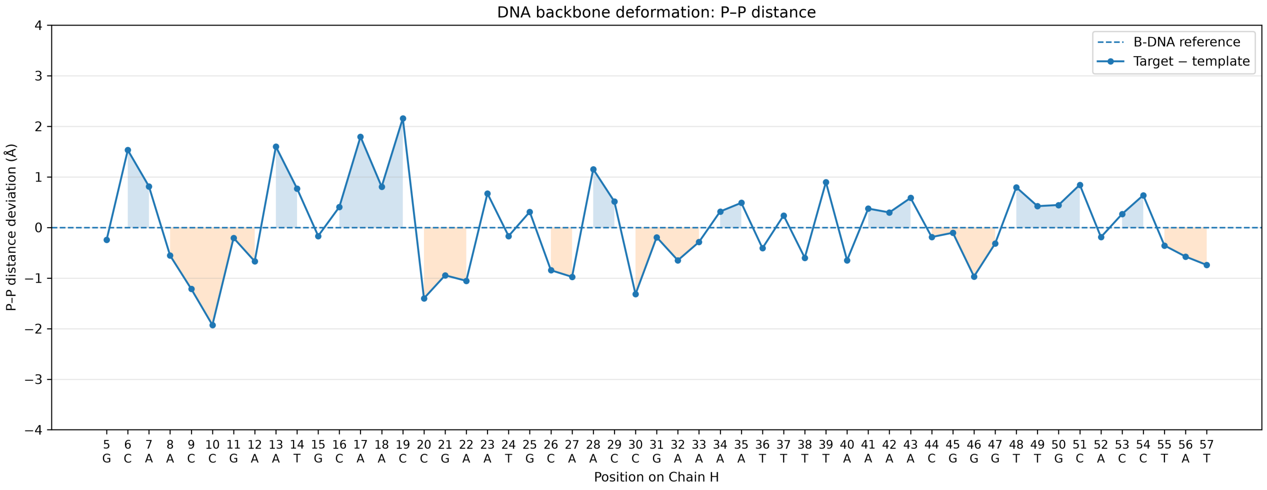


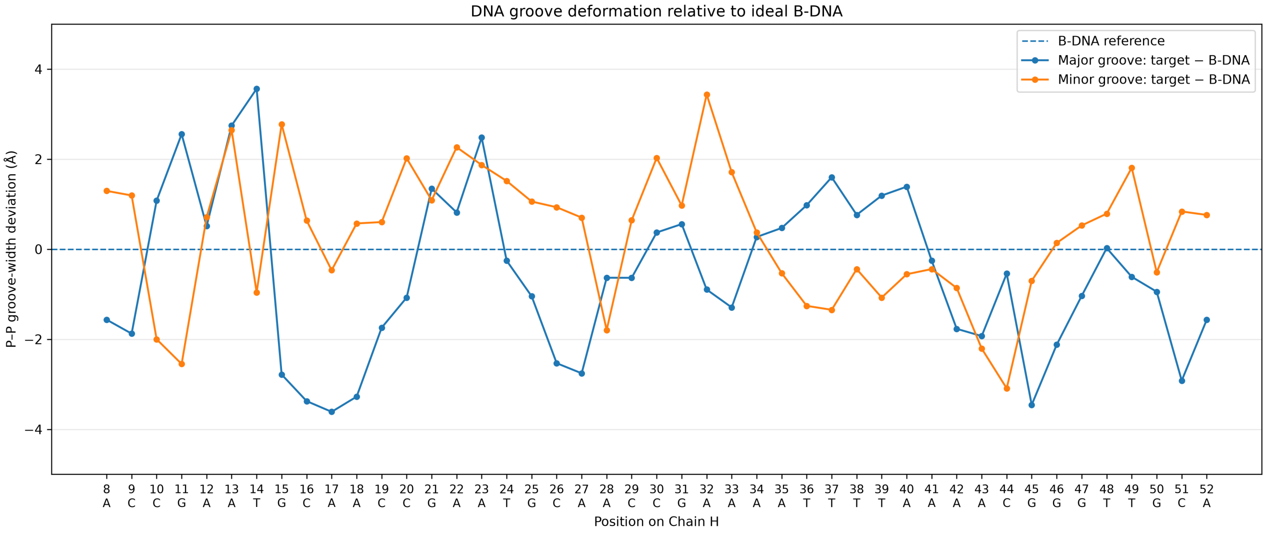


#### PriRep5-DNA-distorted

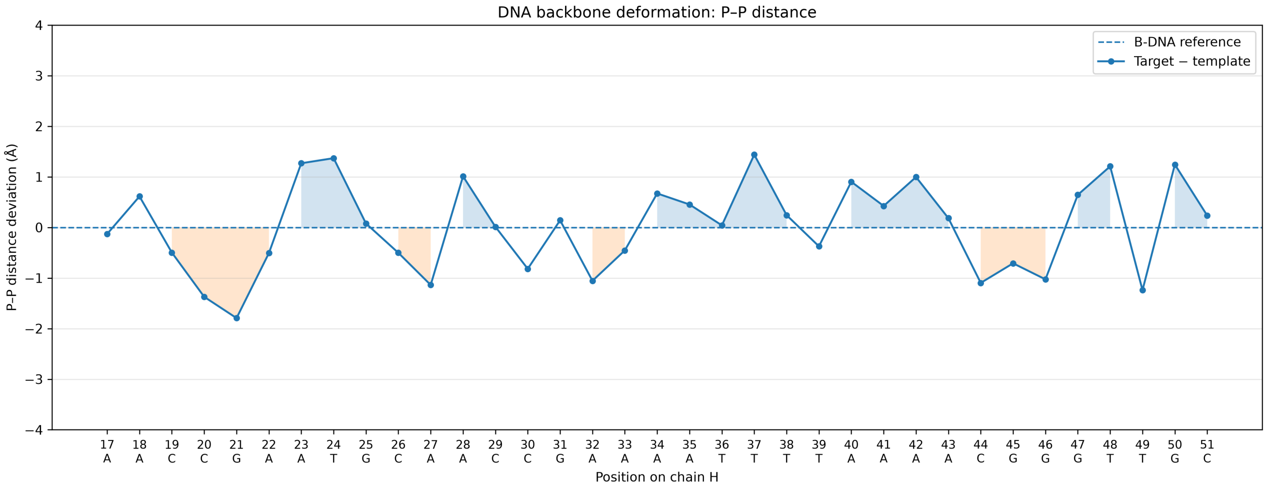


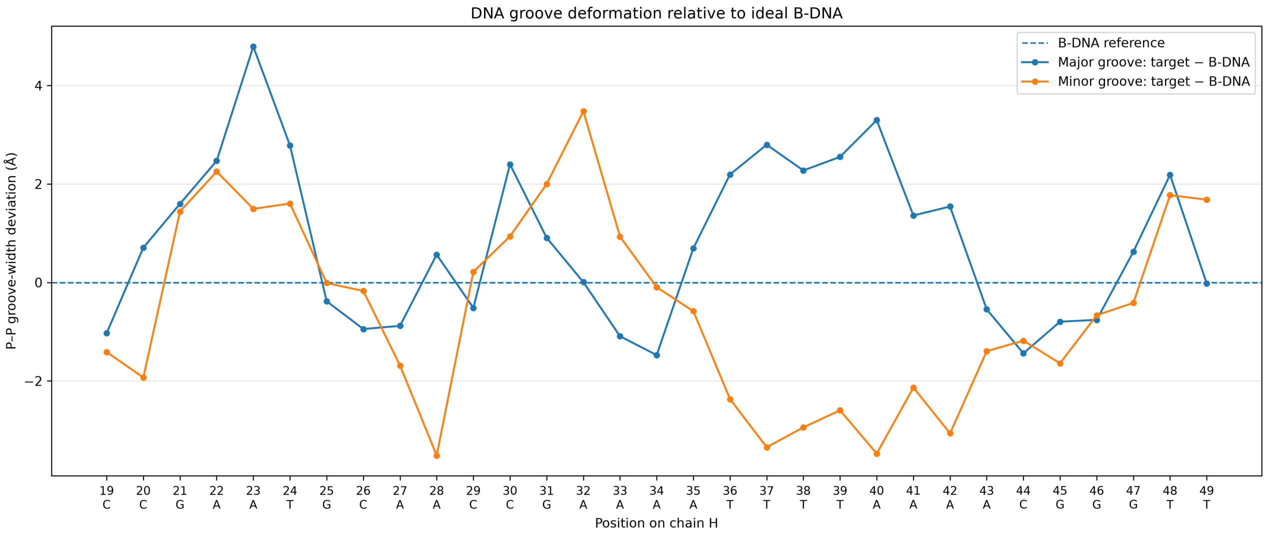

### Supplementary Figure 6

Supp Figure DNA movements multibody intermediates

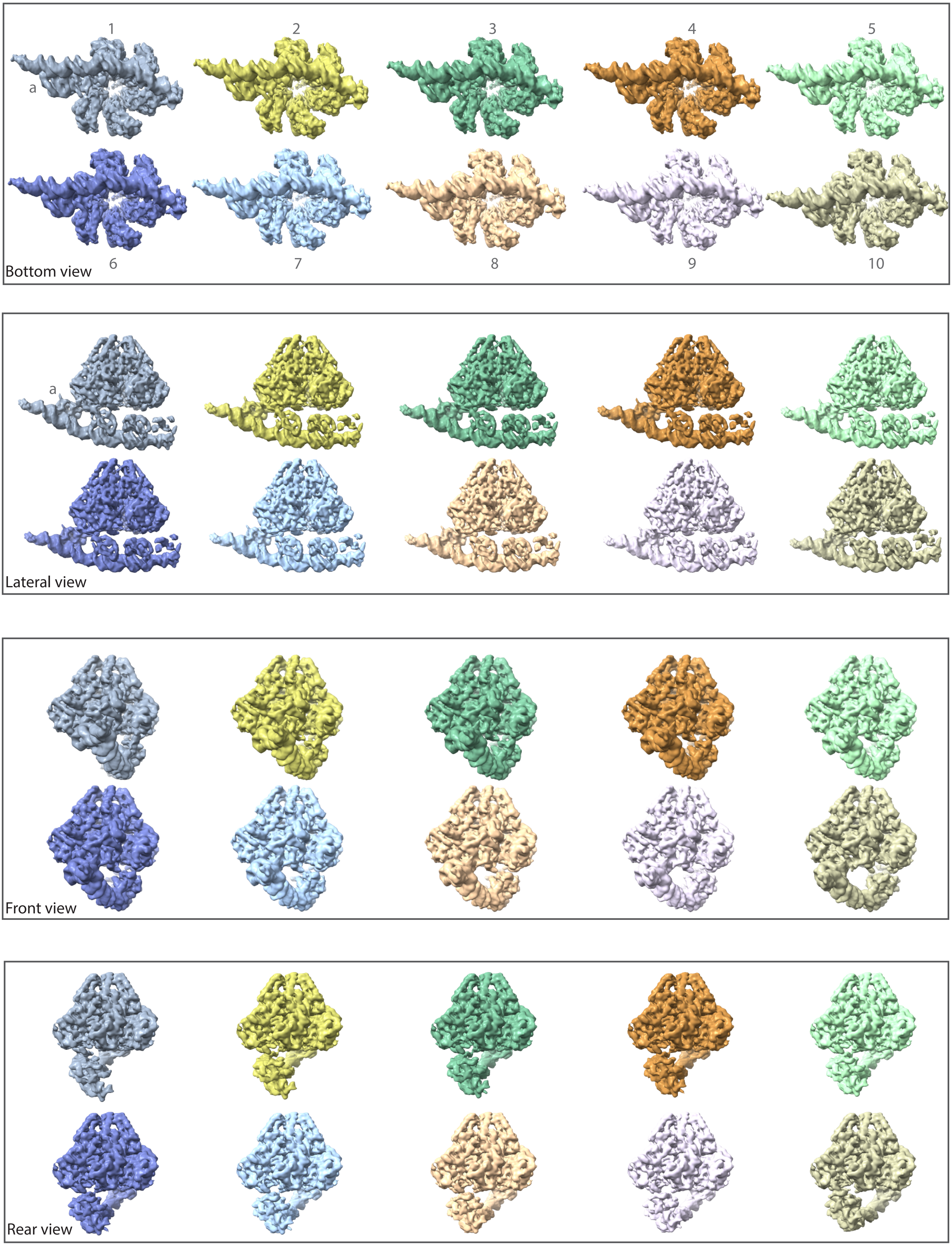

### Supplementary Figure 7

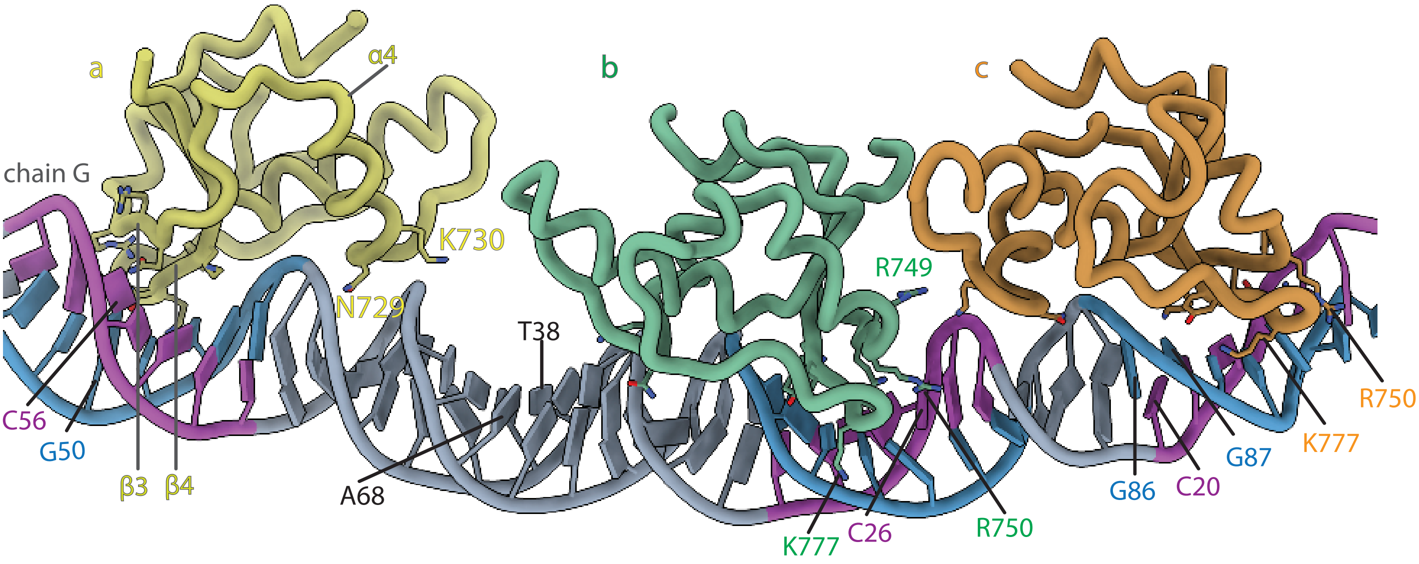

### Supplementary Figure 8

Supplementary figure - PriRep5-DNA-distorted Model to map fit

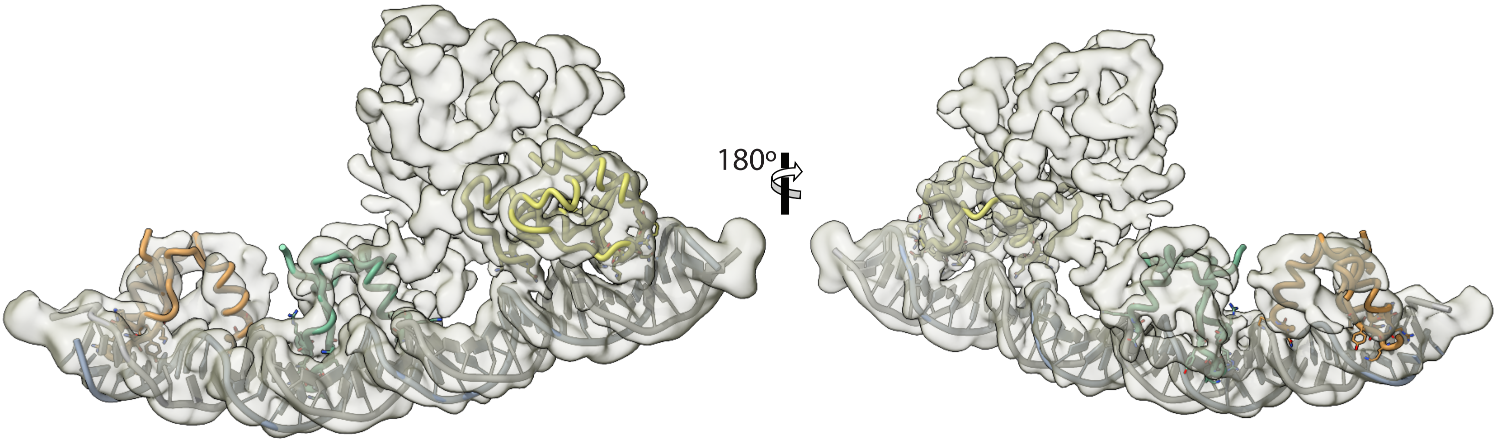
