## Supplementary Figure 2 -FSC for "Structural basis of self-loading by replicative helicases"

PriRep5-4CTDs

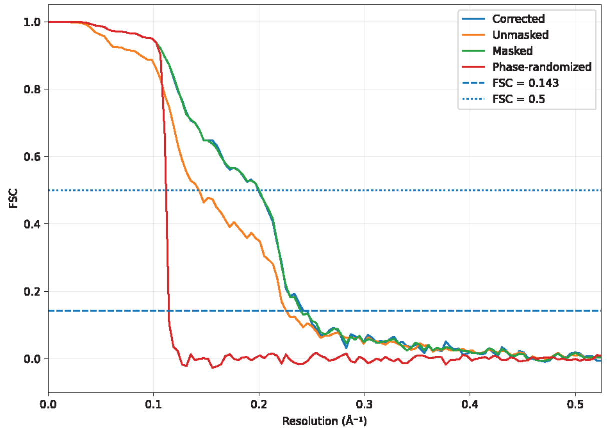


FSC Consensus Map at 4.2 Å resolution [EMD-59247]

PriRep5-4CTDs

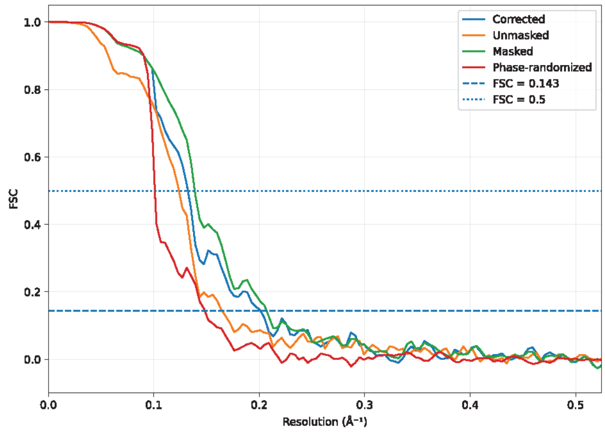


FSC Focused Map at 4.9 Å resolution [EMD-59248]

PriRep5-DNA-distorted

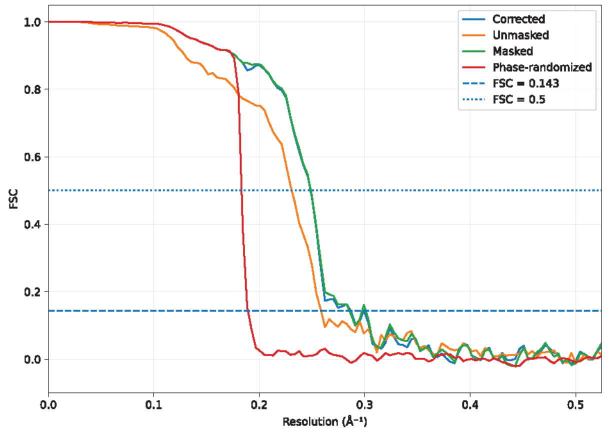


FSC Focused Helicase Map at 3.5 Å [EMD-58827]

PriRep5-DNA-distorted

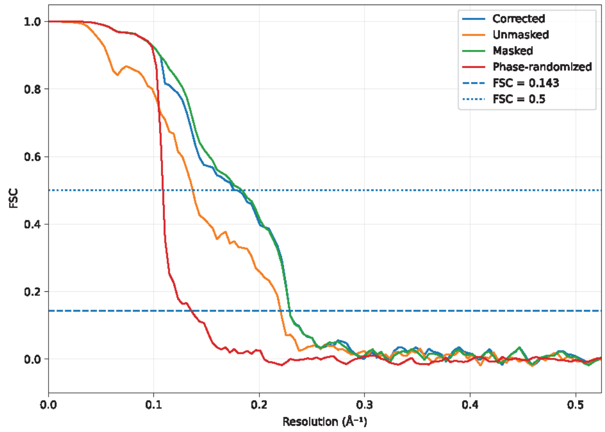


FSC Focused CTDs-DNA distorted Map at 4.4 Å [EMD-58829]

PriRep5-4CTDs

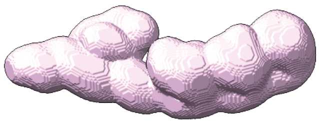

PriRep5-DNA-distorted

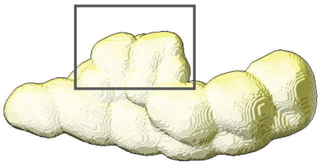

Consensus Map - PriRep5-4CTDs

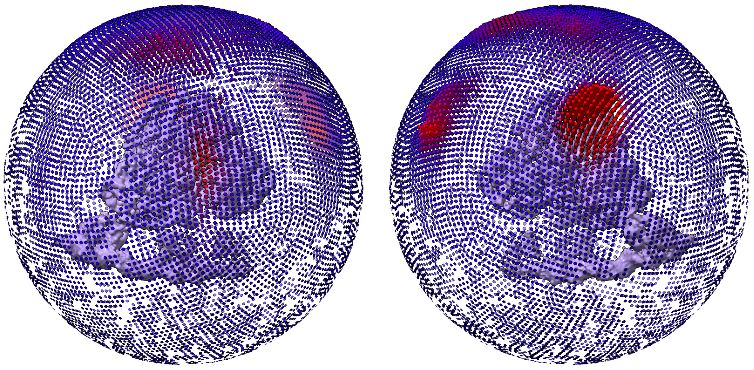

Consensus Map - PriRep5-DNA-distorted

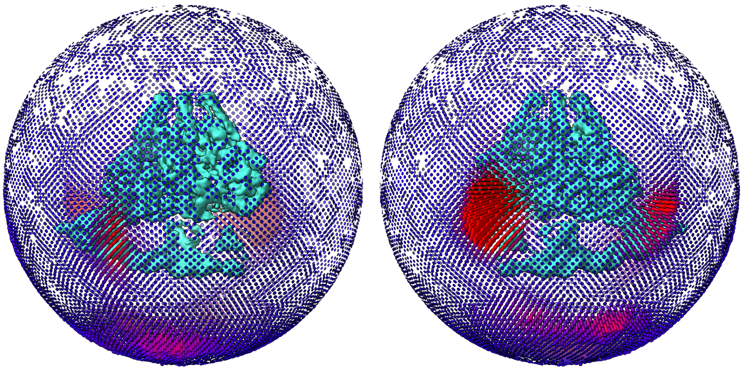
