## Supplementary Figure 4 for "Structural basis of self-loading by replicative helicases"

a

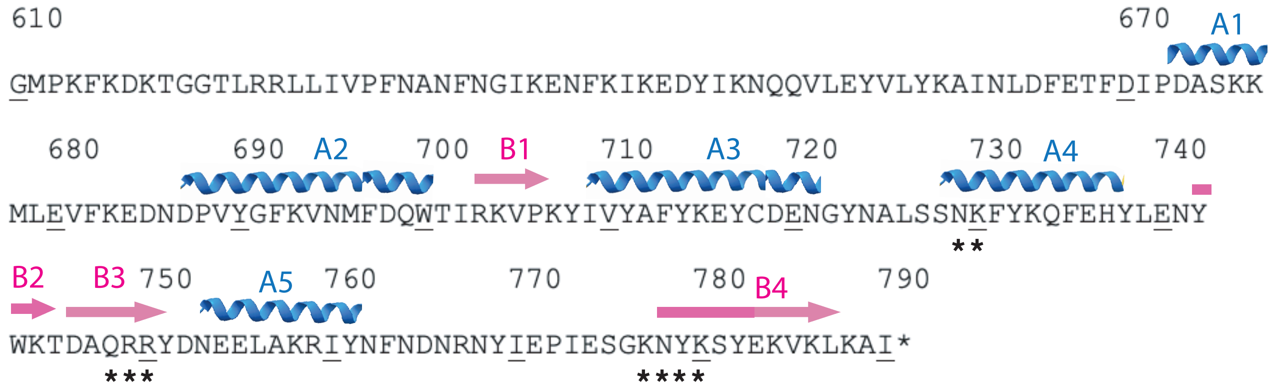


b

| | upstream | downstream |
|---|---|---|
| SaPI5 (80/76.4/79.4 %) | tgcaacc gaaaattttaaaacg ggttgca | tgcaac tggaaaatttaaaacga gttgca |
| SaPIBov1 (86.6/67.6/80.9 %) | gtaccc ataaataaatgtataac gggtac | gtaccc gaagtaaaagaatcgctg gggac |
| SaPI2 (64.7/76.4/80 %) | agttccc gcgaaaaacatacaaac gggaact | agttccc gataagtattttaagtc gggaatt |
| SaPI1 (71.4/68.7/74.6 %) | catctt atttttaggacagt aagaca | tgtctt attcgaattgcaattc aagaca |
